# An engineered IdeS variant with enhanced activity and performance for IgG degradation

**DOI:** 10.64898/2026.06.26.734701

**Authors:** Keyi Zhang, Wenhao Ma, Zhijie Wu, Zipei Ren, Congcong Chen, Yan Xia, Dan He, Ziting Yu, Hongze Niu, Jing Qin, Pan Gao, Wenyuan Yang, Yangguang Dai, Xiu Li, Zheyue Dong, Yue Wang, Xiaoyan Dong, Cheng Chen, Xiaobing Wu

## Abstract

IgG-degrading enzymes have emerged as innovative therapeutic agents for treating conditions driven by pathogenic antibodies. Here, we used structure-guided rational design to engineer IdeS^M33^, a double mutant (K167R/D226E) of the IgG-specific bacterial protease IdeS from *Streptococcus pyogenes*, with improved catalytic efficiency. Biolayer interferometry revealed a fourfold increase in binding affinity relative to wild-type IdeS (IdeS^WT^). This enhancement is likely attributable to mutations that strengthen hydrogen bonding at the enzyme–IgG Fc interface. In vitro, IdeS^M33^ has higher performance than IdeS^WT^ in cleaving serum IgG. In vivo studies in rabbits demonstrated that IdeS^M33^ effectively depleted circulating IgG and showed better performance at a dose of 0.005 mg/kg than the IdeS^WT^. Although doses greater than 0.2 mg/kg demonstrated higher plasma concentrations of IdeS and a larger AUC₀₋last, they did not show a significant enhancement in the pharmacodynamics of IgG degradation. Importantly, a single dose of IdeS^M33^ (0.2 mg/kg) potently degraded binding and neutralizing antibodies against AAV9 within 1-2 days and restored hepatic AAV9 transduction in pre-immunized animals. Together, these findings highlight IdeS^M33^ as a potent and safe engineered enzyme with therapeutic potential for autoimmune disorders, transplant rejection, and overcoming pre-existing humoral immunity in gene therapy.

## Introduction

Effective modulation of the immune system remains a paramount challenge across multiple fields of modern medicine, including the management of autoimmune diseases, the prevention of allograft rejection in organ transplantation, the mitigation of anti-drug antibody responses against biologic therapeutics, and the neutrpalization of pre-existing immunity to viral vectors in gene therapy.^1–5^ Each of these scenarios is critically dependent on the humoral immune response, where immunoglobulin G (IgG) antibodies serve as the primary effector molecules.^1–3^ Conventional strategies to control pathogenic or inconvenient antibodies—such as plasmapheresis, immunosuppressive drugs, or B-cell depletion—are often non-selective, associated with significant side effects, or offer only transient efficacy.^1–3^ The advent of highly specific bacterial enzymes capable of selectively cleaving IgG has revolutionized the clinical toolkit for antibody depletion.^1–5^ Among these, the IgG-degrading enzyme of *Streptococcus pyogenes* (IdeS) has emerged as a transformative agent, offering a precise, rapid, and potent method to eliminate circulating IgG, thereby changing the management paradigm for these previously intractable problems.^1–6^

IdeS, a 35 kDa cysteine protease, was first identified in the human pathogen *Streptococcus pyogenes*.^7^ It exhibits exquisite specificity, cleaving human IgG after the Gly236 residue, generating one F(ab’)2 fragment and two homodimeric Fc fragments.^7^ This proteolytic event irreversibly abolishes the antibody’s effector functions, such as complement-dependent cytotoxicity and antibody-dependent cellular cytotoxicity, while also preventing Fcγ receptor-mediated signaling.^7,8^ The clinical potential of this bacterial “molecular scalpel” was swiftly recognized.^1–6^ Its recombinant form, imlifidase, received conditional marketing authorization in the European Union in 2020 (and later in the US) for the desensitization of highly sensitized adult kidney transplant patients.^9,10^ Its application has since expanded into clinical trials for treating antibody-mediated autoimmune disorders, such as anti-glomerular basement membrane disease and ANCA-associated vasculitis, and is being actively explored as a pre-treatment to enable effective gene therapy with adeno-associated virus (AAV) vectors in seropositive individuals.^11,12^

The quest for improved IgG-degrading enzymes has spurred global efforts in protein engineering, exploring diversity across species and employing various molecular techniques.^13,14^ Beyond the streptococcal IdeS, homologous enzymes have been identified in other bacteria, such as *Streptococcus equi* (IdeE) and *Streptococcus canis*, each with distinct IgG specificities and catalytic properties.^15,16^ Protein engineering strategies to enhance these enzymes have included: 1) Direct evolution to improve catalytic efficiency or stability; 2) Humanization to reduce immunogenicity; 3) Fusion protein strategies, such as creating Fc fusions to prolong serum half-life; and 4) Rational design based on structural insights to modify substrate affinity or alter specificity towards different IgG subclasses.^5,8,17–19^ The overarching goal is to engineer next-generation variants with superior pharmacokinetics, broader or altered subclass specificity, reduced immunogenicity, or enhanced activity in complex biological matrices like human serum.

This study is grounded in the paradigm of structure-guided rational design to achieve functional enhancement. While wild-type IdeS is highly efficacious, opportunities for improvement exist, particularly in enhancing its affinity for IgG to ensure complete depletion even at low enzyme concentrations.^19^ Through computational analysis of the co-crystal structure of IdeS bound to its IgG Fc substrate, we performed mutagenesis focused on optimizing interfacial electrostatics and hydrogen-bonding networks. This effort led to the identification of a double mutant, D226E and K167R, which displayed enhanced in vitro and in vivo performance in IgG degradation.

## Results

### Rational design of IdeS^M33^

To improve the activity of IdeS MG50 (IdeS^WT^), we first reconstructed the IdeS MG50–IgG-Fc complex structure using the crystal structure of the IdeS MG83–IgG-Fc complex (PDB ID: 8A47) as a template with PyRosetta (Fig. 1A). Ligplot analysis of the hydrogen-bond networks between IdeS MG50 (chain C) and IgG-Fc (chains A and B) identified key interacting residues. Specifically, the B-C interface involved E132, K167, R185, T189, and R204, while the A-C interface involved E66, D67, K84, A94, D226, K228, V258, R259, A310, and G319 (Fig. 1B & Supplementary Fig1. A - B).

**Figure 1.**
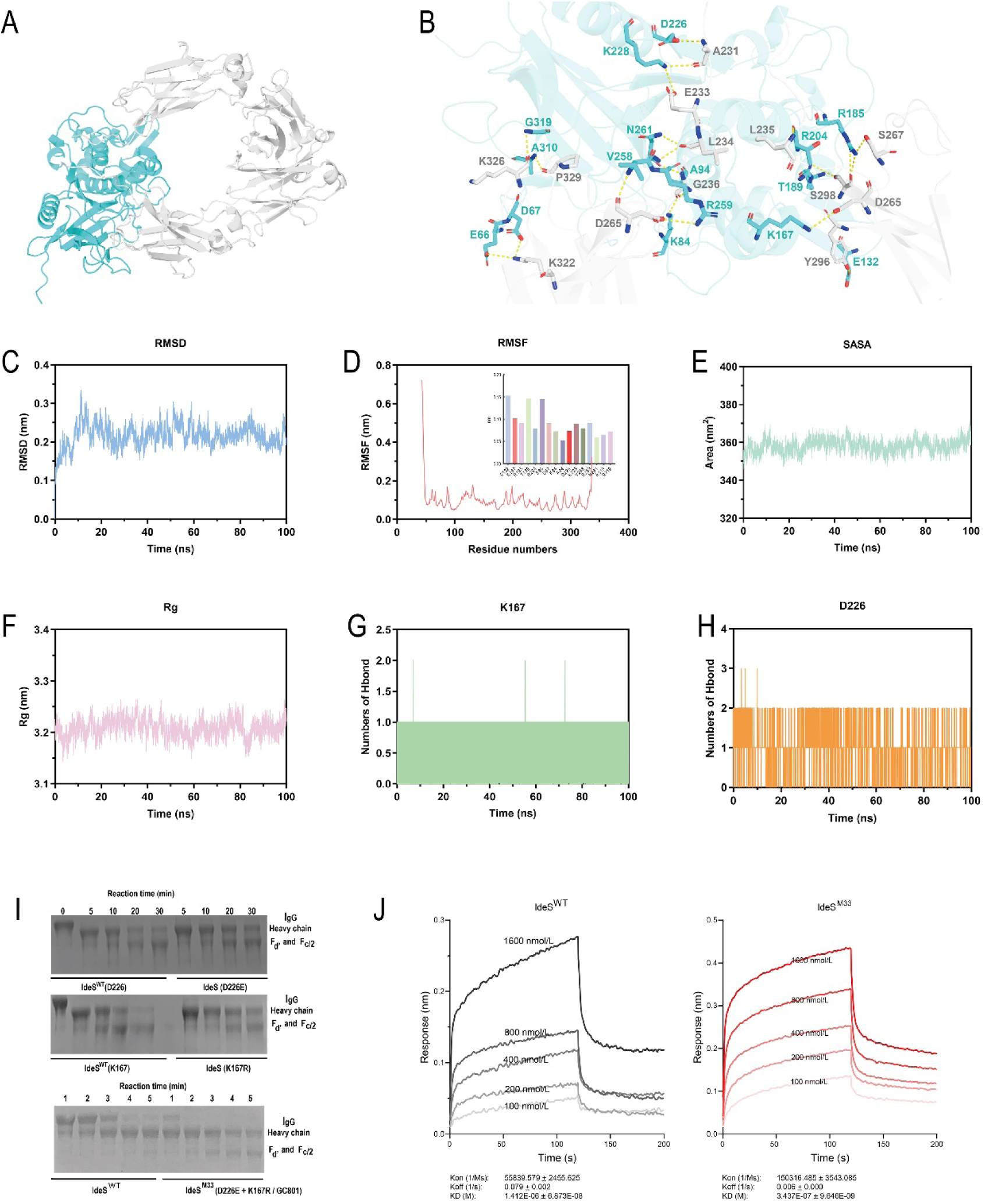
Rational design of IdeS^M33^ and functional validation. A. The structure of the IdeS MG50–IgG-Fc complex was modeled using the crystal structure of the IdeS MG83–IgG-Fc complex (PDB ID: 8A47) as a template with pyRosetta. B. Ligplot analysis of the hydrogen-bond network between IdeS MG50 (chain C) and IgG-Fc (chains A and B) identified key interacting residues. The B-C interface involved E132, K167, R185, T189, and R204, while the A-C interface involved E66, D67, K84, A94, D226, K228, V258, R259, A310, and G319. C. Root mean square deviation (RMSD) analysis. D. Root mean square luctuation (RMSF) analysis. E. Solvent-accessible surface area (SASA) trajectory. F. The radius of gyration (Rg) values. G. Hydrogen-bond dynamics analysis focused on residue K167. H. Hydrogen-bond dynamics analysis focused on residue D226. I. A. Analysis of IgG cleavage efficiency of individual mutants and double mutant (IdeS^M33^) compared to IdeS^WT^. J. Biolayer interferometry measurement of *k*_on_, *k*_off_, and *K*_D_ for the interaction of IdeS^WT^ and IdeS^M33^ with IgG.

To evaluate the dynamic behavior of interface residues in solution, we performed a 100 ns MD simulation of the IdeS MG50–IgG-Fc complex. RMSD analysis (Fig. 1C) indicated that the system reached equilibrium after an initial ∼20 ns equilibration phase, with the overall RMSD remaining below 0.3 nm, suggesting a stable global conformation. Consistent with this, RMSF analysis (Fig. 1D) showed that interface residues were located in low-fluctuation (rigid) regions, favoring binding stability; most core regions also exhibited low RMSF values, indicating a robust structural scaffold. Furthermore, the SASA trajectory (Fig. 1E) displayed only minor fluctuations, implying no significant change in solvent accessibility, while the radius of gyration (Rg) fluctuated smoothly (variation < 0.1 nm) (Fig. 1F), confirming the absence of tertiary structural expansion or contraction.

Hydrogen-bond dynamics analysis (Supplementary Fig. 2) revealed distinct behaviors at the two interfaces. At the B-C interface, R185, T189, R204, and E132 formed extremely robust networks, exhibiting occupancies >99% and maximum continuous absence windows of ≤10 ps; notably, R185 formed an average of 7.5 hydrogen bonds. In marked contrast, K167 exhibited poor hydrogen-bond maintenance, with an average of only 0.71 hydrogen bonds, an occupancy of merely 70.65%, and a maximum continuous absence window of 90 ps (Fig. 1G). Although the RMSF of K167 was low (0.102 nm), indicating inherent rigidity, its weak interaction likely stems from suboptimal side-chain geometry rather than flexibility, thus identifying it as a prime candidate for engineering. Similarly, at the A-C interface, while most residues formed stable hydrogen bonds, D226 and R259 were less stable. D226 showed an average of 1.01 hydrogen bonds and an occupancy of 96.38%, whereas R259 exhibited an average of only 0.96 bonds, an occupancy of 85.21%, and a prolonged absence window of 260 ps (Fig.1H). Given that R259 already possesses a flexible guanidinium group, its chemical space for optimization is limited. Conversely, the side chain of D226 contains only a single methylene-linked carboxyl group, leaving room for extension to improve geometric complementarity.

Based on these findings, we established a rational design strategy: for interface residues forming unstable yet optimizable hydrogen bonds, we aimed to extend the side chain and enhance polarity via mutation to reinforce the hydrogen-bond network. Following this rationale, K167 was mutated to arginine (K167R) to introduce a longer side chain with a guanidinium group capable of forming multiple hydrogen bonds. D226 was mutated to glutamate (D226E) to add one methylene group, allowing the carboxyl group to more readily access the Fc surface without altering charge character. Consequently, we designed two single-point mutants, K167R and D226E, as well as a double mutant M33 (K167R/D226E) combining both substitutions (which was designated as IdeS^M33^ or GC801).

### IdeS^M33^ exhibited the optimal cleavage efficiency

The in vitro IgG cleavage assay, performed with the expressed and purified mutants, demonstrated that: at identical concentrations of purified IgG and IdeS, the D226E and K167R single mutants exhibited cleavage time courses similar to that of IdeS^WT^, showing prolonged cleavage durations (Figure 1I). In contrast, the D226E + K167R double mutant completed cleavage of the target IgG within 5 min (Figure 1I).

### IdeS^M33^ exhibited the optimal IgG affinity

Biolayer interferometry analysis revealed that both IdeS^WT^ and IdeS^M33^ exhibit extremely rapid IgG-binding kinetics, as evidenced by a nearly imperceptible baseline phase. Notably, the response curve of 400 nM IdeS^M33^ was comparable to that of 1600 nM IdeS^WT^. The association rate constant (*k*_on_) of IdeS^M33^ (150,316.485 ± 3,543.085 M⁻¹s⁻¹) was higher than that of WT IdeS, while its dissociation rate constant (*k*_off_) (0.006 ± 0.000 s⁻¹) was lower. Consequently, the equilibrium dissociation constant (*K*_D_) of IdeS^M33^ (3.437 × 10⁻⁷ ± 9.646 × 10⁻⁹ M) was lower than that of IdeS^WT^. Collectively, these three parameters indicate that, compared to IdeS^WT^, IdeS^M33^ possesses a higher IgG association rate, a lower dissociation rate, and enhanced binding affinity (Figure 1J).

### Structural characterization of IdeS^M33^

The crystal structure of IdeS^M33^ (amino acids 47-339. PDB ID 26PT, Extended PDB ID pdb_000026PT) was determined at 1.91 Å resolution using X-ray diffraction (SSRF BL02U1, λ = 0.979183 Å) in space group *P*2_1_2_1_2_1_, with unit cell parameters a=64.3, b=87.7, c=56.4 Å. Data collection achieved high completeness (99.0% for overall data, 88.0% for highest-resolution shell data, the same below) and redundancy (11.6, 6.8), with low *R*_merge_ (5.6%, 63.5%) and excellent CC_1/2_(0.999). Refinement converged with R_work_/R_free_ of 18.8%/21.1% (28.9%/32.4% at highest-resolution shell) using 25,064 reflections (2,197 in highest-resolution shell), and 2,471 non-hydrogen atoms (2,333 protein, 138 solvent). The model exhibits excellent stereochemistry: r.m.s. deviations of 0.006 Å in bond lengths and 0.83° in bond angles, with 97.6% of residues in favored Ramachandran regions and 0.0% outliers. These metrics indicate a well-ordered, high-quality structure suitable for detailed mechanistic and functional analysis (Supplementary Table 1).

Although several crystal structures of IdeS have been reported to date, co-crystallizing IdeS with IgG remains highly challenging. This challenge stems from IdeS’s efficient and specific IgG - cleaving activity, which has precluded the direct structural determination of catalytically active wild - type IdeS in complex with IgG. To address this limitation, we determined the crystal structure of the IdeS^M33^ mutant and performed structural alignment with previously reported wild type IdeS structures. This alignment enabled a comparison of their overall conformations and key functional regions, providing a structural basis for further elucidating the conformational features and functional alterations of IdeS^M33^.

To define the structural consequences of the M33 mutations, we superimposed the IdeS^M33^ structure onto the wild type IdeS structure (PDB: 1Y08). The overall protein fold remained largely unaltered, indicating that the mutations are well - tolerated at the global structural level. In contrast, several local regions exhibited distinct conformational differences (Figure 2A).

**Figure. 2.**
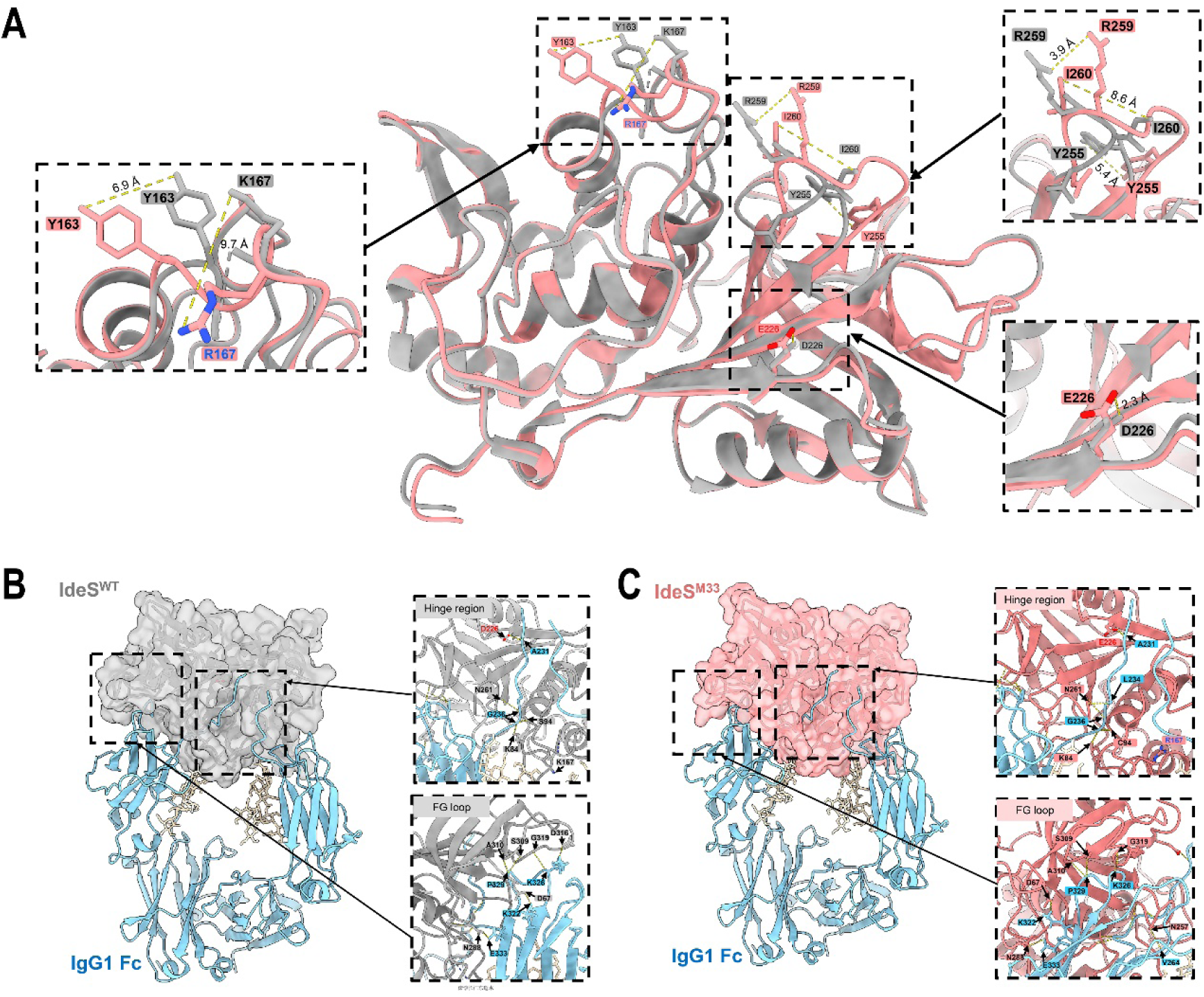
Structural comparison of IdeS^M33^–IgG1 Fc and IdeS^WT^–IgG1 Fc complex models. A. Structural superposition of IdeS^WT^ (gray) and IdeS^M33^ (pink). Mutated residues are highlighted as sticks, and yellow dashed lines indicate measured Cα positional displacements between the aligned structures. Insets show magnified views of key interface regions: left inset highlights Y163–R167 interaction (6.9 Å distance); top-right inset shows R259–Y255 interaction (5.8 Å); bottom-right inset displays E226–D226 interaction (3.3 Å). B and C. Complex models were generated in PyMOL using the IdeS–IgG1 Fc complex structure (PDB ID: 8A47) as a template, in which the IdeS component was replaced with either the IdeS^M33^crystal structure or the IdeS^WT^ structure (PDB ID: 1Y08). Left panels show full complex views: IdeS^WT^ in gray (B), IdeS^M33^ in pink (C); IgG1 Fc in light blue. Right panels provide close-up views of the hinge region and FG loop. Yellow dashed lines denote predicted hydrogen bonds, with participating residues labeled.

The most prominent change occurred near residue 167: the K167R substitution induced a pronounced repositioning of the side chain, with R167 shifted by ∼9.7 Å relative to K167 in the wild type structure. This local rearrangement was accompanied by a 6.9 Å shift of the neighboring Y163 residue, implying that the mutation reshapes the surrounding side - chain environment (rather than acting as an isolated substitution) (Figure 2A).

Additional deviations were observed in the region containing Y255, R259, and I260. Although these residues were not directly mutated, they exhibited shifts of ∼5.4 Å, 3.9 Å, and 8.6 Å, respectively, in the aligned structures. Thus, the structural effects of IdeS^M33^ are not confined to the mutation sites but extend to nearby surface or loop regions (Figure 2A).

In contrast, the D226E mutation caused only a modest displacement (∼2.3 Å). This limited change is consistent with the conservative nature of the substitution (i.e., aspartate to glutamate, both negatively charged), although the longer glutamate side chain may still influence local packing or side - chain contacts. Overall, the alignment analysis reveals that IdeS^M33^ preserves the wild type IdeS scaffold while introducing discrete local conformational changes—most notably around R167 and the Y255–I260 region (Figure 2A).

To compare potential differences in IgG1 Fc recognition between IdeSM33and IdeSWT, we generated two complex models in PyMOL using the IdeS–IgG1 Fc complex structure (PDB: 8A47) as a template. In these models, the IdeS molecule from the template was replaced either with the wild type IdeS structure (PDB: 1Y08) or with the crystal structure of IdeS^M33^ determined in this study.

Local interface comparison revealed differences in the predicted hydrogen - bonding networks around the hinge region and the FG loop:

Hinge region: In the IdeS^WT^ model, hydrogen bonds primarily involved D226, N261, S94, K84, and Fc residues (e.g., A231, G236) (Figure 2B). In the IdeS^M33^ model, the hydrogen - bonding network in this region was partially altered, with an additional interaction observed between N261 and Fc L234 (Figure 2C).

FG loop: Both models formed hydrogen bonds near Fc residues P329, K326, K322, and E333. However, the IdeS^M33^ model exhibited a hydrogen bond between N257 and Fc V264 (Figure 2C), whereas the IdeS^WT^ model contained an additional hydrogen bond between D316 and Fc K326 (Figure 2B).

These results suggest that IdeS^M33^ retains the overall spatial mode for recognizing the Fc hinge–FG loop region but differs from IdeS^WT^ in its local hydrogen - bonding network. This difference may alter the local stabilization of the IgG1 Fc interface.

### IdeS^M33^ exhibits higher cleavage activity for human or rabbit serum IgG

25 μL of serum was treated with 5 μL of IdeS (2 μg/μL) at 37 °C for 1 to 240 min. The relative abundance of IgG heavy chain was visualized and quantified. Both IdeS^WT^ and the IdeS^M33^ began to cleave serum IgG within 5 min and exhibited higher cleavage activity toward human (Figure 3A and B) and rabbit (Figure 3G and H) serum IgG than toward cynomolgus monkey (Figure 3C and D) and canine serum IgG (Figure 3E and F). Cleavage of human and rabbit serum IgG by both enzymes was sustained for almost 240 min, with the relative abundance of IgG heavy chain decreasing over time. One-way ANOVA of the 7 time points for human serum IgG cleavage by IdeS^WT^ and IdeS^M33^ yielded *P*-values of 0.0217 and 0.0013, respectively. For rabbit serum IgG cleavage, the *P*-values for the 7 time points were 0.0065 and 0.0028, respectively. After 240 min, the relative heavy-chain abundance of human IgG for IdeS^WT^ decreased to 53.55 ± 5.56%, and for IdeS^M33^ to 18.97 ± 7.77%. For rabbit IgG, the values decreased to 23.83 ± 6.58% (IdeS^WT^) and 5.74 ± 1.76% (IdeS^M33^). The AUC for human serum IgG cleavage by IdeS^WT^ and IdeS^M33^ over 240 min was 448.0 ± 16.49 and 350.5 ± 27.64, respectively. For rabbit serum IgG cleavage, the AUC values were 358.7 ± 45.05 (IdeS^WT^) and 232.8 ± 16.67 (IdeS^M33^). IdeS^M33^ demonstrated superior IgG-cleavage capability compared to IdeS^WT^ (Figure 3). Based on the above experimental evidence, we used rabbits as the research subjects in the subsequent in vivo enzymatic activity studies.

**Figure 3.**
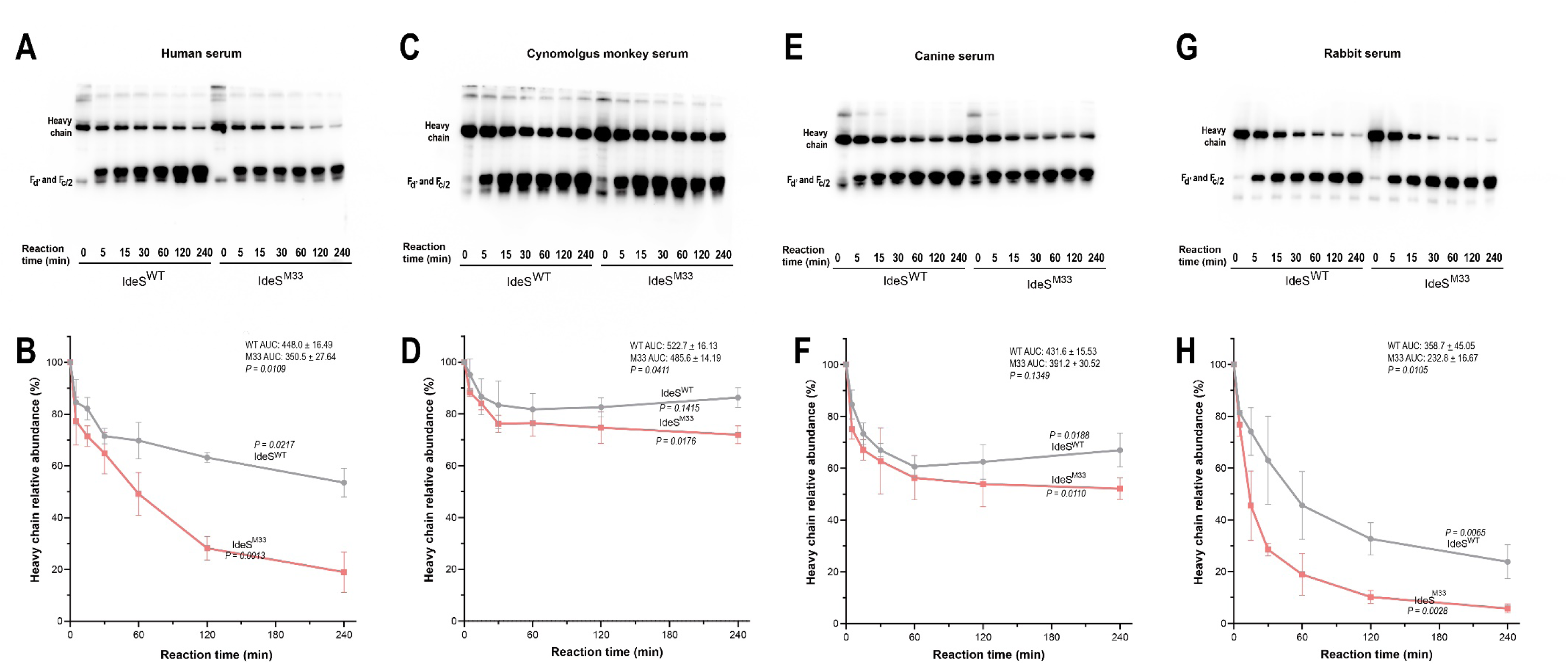
In vitro cleavage of serum IgG from different species by IdeS^WT^ and IdeS^M33^. 25 μL of serum from human, cynomolgus monkey, canine, and rabbit was treated with 5 μL of IdeS (2 μg/μL) at 37 °C for 1 to 240 min. Cleavage efficiency was visualized and quantified by SDS-PAGE and Western blotting. Each serum sample was tested in triplicate for each enzyme. The area under the curve (AUC) was calculated as the product of the reaction time (from 0 to 240 min) and the relative abundance of the heavy chain (the *P*-value represents the result of a t-test for the difference in AUC). The *P* values indicated on the curves were derived from one-way ANOVA testing of the relative abundance of the heavy chain at the seven time points.

### In vivo IgG-degrading capability of IdeS^M33^ versus IdeS^WT^ in rabbits

Following intravenous administration of IdeS^M33^ and IdeS^WT^ at doses of 0.005, 0.01, and 0.2 mg/kg, serum IgG levels were measured on days 1, 2, 3, 7, and 14. As shown in Figure 4, serum IgG levels in rabbits decreased rapidly, reaching their nadir on days 1-2 post-administration. In rabbits treated with 0.005, 0.01, and 0.2 mg/kg of IdeS^M33^ and IdeS^WT^, IgG levels declined to 20.7 ± 2.2% and 25.7 ± 2.6%, 13.6 ± 3.9% and 13.2 ± 0.2%, and 9.5 ± 1.6% and 10.9 ± 2.6% (relative to day0), respectively. IgG levels began to recover from day 3 onward. By day 14, IgG levels in the respective treatment groups had recovered to 59.5 ± 4.1% and 83.2 ± 3.4%, 52.7 ± 15.7% and 69.8 ± 19.5%, and 48.0 ± 11.2% and 74.0 ± 40.3% of pre-dose levels.

**Figure 4.**
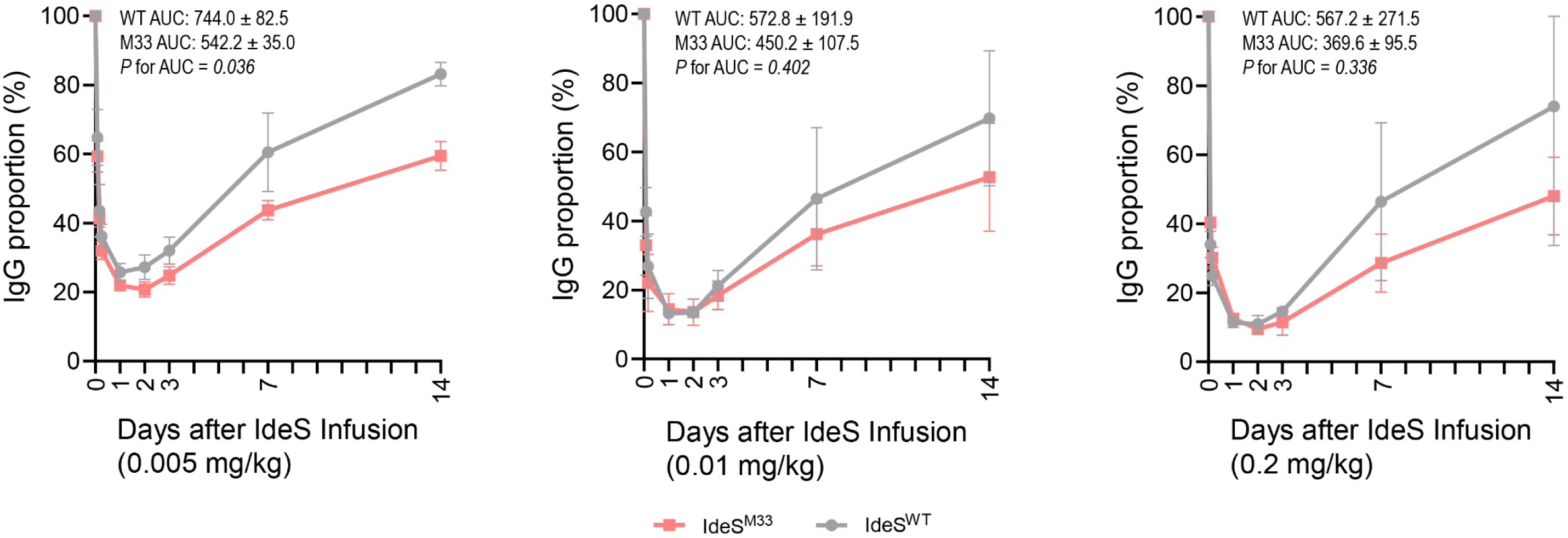
A comparison of in vivo IgG-degrading capability. In vivo IgG-degrading capability of IdeS^WT^ versus IdeS^M33^ in rabbits. Following intravenous administration of IdeS at doses of 0.005, 0.01, and 0.2 mg/kg (N=3 for each group), serum IgG levels were measured on days 1, 2, 3, 7, and 14. The values are presented as percentages relative to the IgG level on day 0 (pre-dose).

After IdeS infusion, for both the IgG nadir and the recovery percentage at day 14, IdeS^M33^-treated groups showed lower values than IdeS^WT^-treated groups in most cases. IdeS^M33^ demonstrated smaller AUC values across all three dose groups. Specifically, in the 0.005 mg/kg dose group, the AUC for IdeS^M33^ was 542.2 ± 35.0, which was significantly smaller than the AUC for IdeS^WT^ (744.0 ± 82.5; P = 0.036). These results indicate that IdeS^M33^ possesses superior IgG-degrading capability, particularly at a dosage of 0.005 mg/kg, compared to IdeS^WT^.

### Pharmacokinetics and pharmacodynamics of IdeS^M33^

As shown in Figure 5A, the rabbit serum IgG proportion in the 0.2 mg/kg, 0.6 mg/kg, and 1.8 mg/kg groups decreased to 8.59 ± 1.96%, 8.22 ± 2.64%, and 6.87 ± 3.01%, respectively one day post IdeS^M33^ administration. IgG levels then began to rise beyond day 3, reaching 39.81 ± 12.25%, 40.55 ± 15.67%, and 27.27 ± 10.91%, respectively on day 7. Figure 5B showed the pharmacokinetics. The 1.8 mg/kg group achieves highest total drug exposure. Its C_max_ reaches as high as 59,509.6 ng/mL, which is approximately 31 times that of the 0.2 mg/kg group. Its AUC is also the highest (44,195.9), indicating the greatest total cumulative drug amount in the body. However, the 1.8 mg/kg group has the shortest *t*_1/2_ (13.2 hours). Although it achieves the highest peak concentration, it is also cleared most rapidly, resulting in faster drug elimination. The 0.6 mg/kg group achieves the longest duration of action. Its *t*_1/2_ is the longest (32.0 hours), meaning the drug maintains effective concentrations in the body for the longest time. The low-dose (0.2 mg/kg) group showed the lowest AUC and C_max_, yet its *t*_1/2_ could reach 22.5 hours.

**Figure 5.**
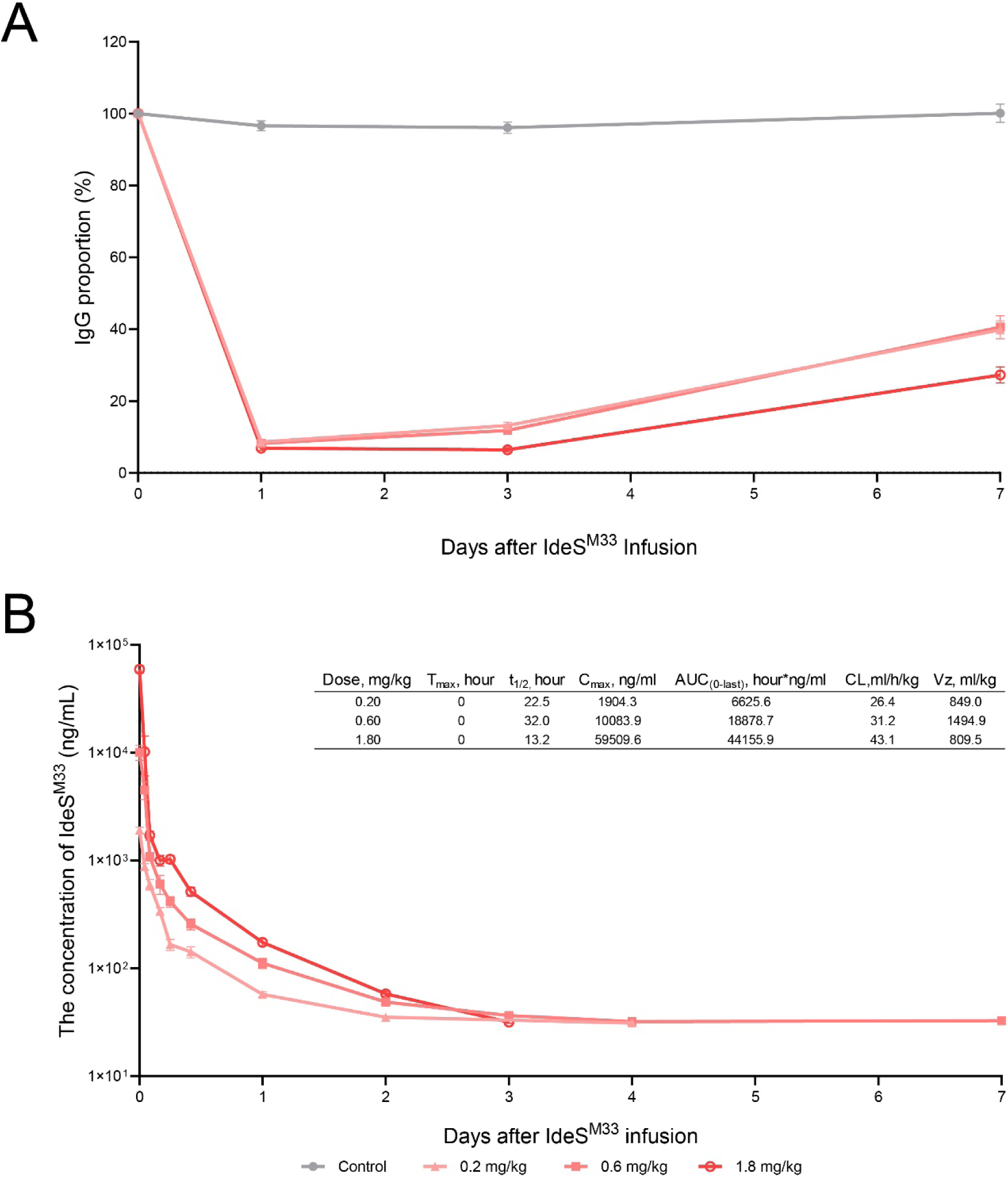
Pharmacokinetics and pharmacodynamics of IdeS^M33^ in vivo. A, Pharmacodynamics in rabbits. New Zealand white rabbits were divided into four groups based on dosage: 0 mg/kg, 0.2 mg/kg, 0.6 mg/kg, and 1.8 mg/kg. IdeS^M33^ was administered intravenously on Day 0. Changes in blood IgG levels were observed until Day 7. B. Pharmacokinetics in rabbits. Blood concentrations of IdeS^M33^ were measured up to Day 7, with six measurements taken on the first day. T_max_, time to maximum plasma concentration; t_₁/₂_, elimination half-life; C_max_, concentration at the maximum; AUC_(0-last)_, area under the plasma concentration-curve from time zero to the last measurable concentration; CL, clearance; V_z_, volume of distribution during the terminal phase.

### IdeS^M33^ demonstrates favorable in vivo safety

As shown in Supplementary Figure 3, intravenous administration of IdeS^M33^ at doses of 0.0, 0.2, 0.6, and 1.8 mg/kg resulted in no statistically significant differences among dose groups in body temperature, body weight, complete blood count, or blood biochemistry parameters—regardless of sex—with the exception of the red blood cell (RBC) count in male rabbits, which was significantly lower in the 1.8 mg/kg group. Rabbits treated with IdeS^M33^ exhibited similar growth trends across all groups (Supplementary Figure 3B and 3C). Although some biochemical parameters displayed increasing or decreasing trends following IdeS^M33^ administration, the patterns of change showed no significant differences among dose groups (Supplementary Figure 3F and 3G). Although rabbit CK levels exceeded the 50–300 U/L reference range, this elevation was not attributed to IdeS^M33^ treatment. The rationale is that the high-dose group (1.8 mg/kg) showed a decreasing trend in CK after injection, whereas CK levels in the control group, though relatively stable before and after administration, also remained above the normal reference range.

### IdeS^M33^ degrades both total and neutralizing antibodies against AAV9

Three weeks after the second immunization, IdeS^M33^ (0.2 mg/kg) or solvent was administered. The levels of neutralizing antibodies and total binding antibodies against AAV9 in the serum were detected before IdeS^M33^ administration and at 24 h, 48 h, and 72 h after administration. As shown in Figure 6A-B, compared to pre-treatment levels, IdeS^M33^ reduced binding antibodies against AAV9 to 1.50 ± 0.87%, at 24 h post-administration. Binding antibody levels then began to rise slowly by 48 h. Similarly, IdeS^M33^ reduced neutralizing antibodies (nAb) against the AAV9 to 1.93 ± 0.76%, at 24 h post-administration. Neutralizing antibody levels likewise began to recover slowly by 48 h.

**Figure 6.**
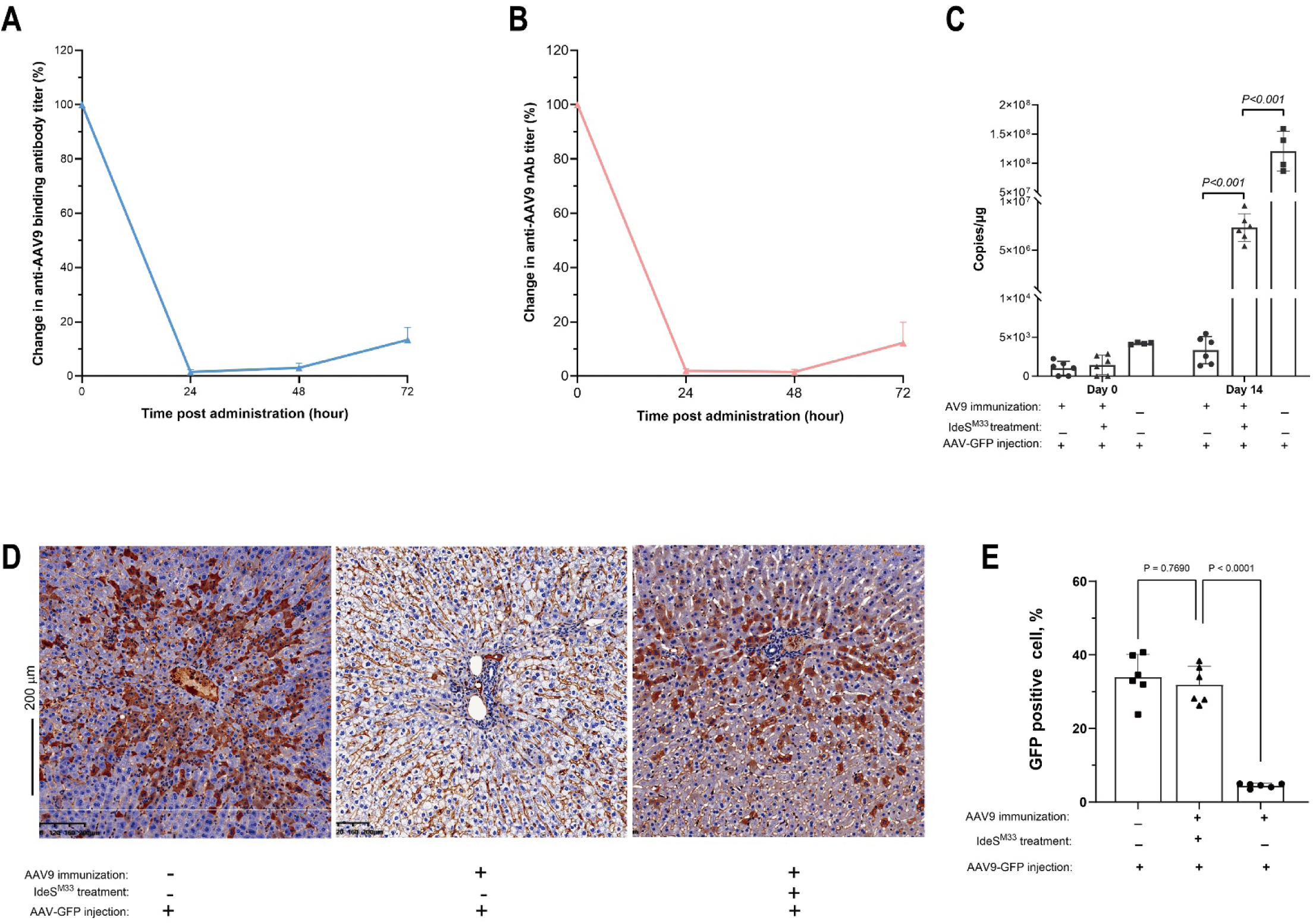
IdeS^M33^ can effectively cleave antibodies against AAV9 and thereby enhance the transduction efficiency. A-B, IdeS^M33^ demonstrated a high efficiency in cleave both total antibodies and neutralizing antibodies against AAV9. C, Rabbits pre-immunized with AAV9 (N=6 per group) were divided into two groups based on whether they received treatment with 0.2 mg/kg IdeS^M33^ or not. One day later (Day 0), all rabbits received an intravenous injection via the tail vein of rAAV9-eGFP (2 × 10¹² vg/animal). Fourteen days later, the copy number of rAAV9-eGFP in liver tissue was measured to quantify the effect of IdeS^M33^ on AAV9 transduction efficiency. D, Fourteen days later, liver histopathology was performed on the rabbits described in panel C to observe GFP expression within the liver across the groups. E, Quantitative analysis of the GFP positive cells in the histopathology images from panel D further supported that IdeS^M33^ can increase the transduction efficiency of AAV9 in the liver of rabbits with pre-existing AAV9 antibodies.

### IdeS^M33^ treatment rescues the transduction efficiency of AAV vectors in rabbits with pre-existing anti-AAV antibodies

Rabbits pre-immunized with AAV9 were either treated with 0.2 mg/kg IdeS^M33^ or left untreated. One day later (designated Day 0), all rabbits received 2 × 10¹² vg/animal of rAAV9-eGFP via tail vein injection. Fourteen days post-injection, the rAAV9-eGFP copy number in liver tissue was measured. The copy number in IdeS^M33^-treated rabbits was (7.33 ± 1.42) × 10⁶. This value was significantly higher than that in AAV9-pre-immunized, but IdeS^M33^ untreated rabbits ((3.34 ± 1.76) × 10³), demonstrating that IdeS^M33^ restored transduction. However, it remained significantly lower than the copy number in rabbits that were not pre-immunized with AAV9 ((1.21 ± 0.34) × 10⁸) (Figure 6C).

Fourteen days later, the results of liver immunohistochemistry were consistent with the quantitative GFP DNA data described above. Rabbits without pre-existing anti-AAV9 immunity exhibited the most extensive GFP (brown) distribution around the portal area of the hepatic lobule. Rabbits with pre-existing anti-AAV9 immunity showed the least GFP distribution. In rabbits with pre-existing anti-AAV9 immunity that were treated with IdeS^M33^, the amount of GFP distribution in the portal areas was intermediate between the two other groups (Figure 6D). Quantification of GFP-positive cells showed that in rabbits with pre-existing anti-AAV9 immunity treated with IdeS^M33^, the GFP positivity rate was 31.87 ± 5.11%. This value was significantly higher than that in rabbits with pre-existing immunity but without IdeS^M33^ treatment (4.45 ± 0.61%), but showed no statistically significant difference compared to the GFP positivity in rabbits without pre-existing anti-AAV9 immunity (33.39 ± 6.13%) (Figure 6E).

## Discussion

In this study, we employed a structure-guided approach to engineer a double mutant, IdeS^M33^ (K167R/D226E), which exhibits significantly enhanced IgG-binding affinity and catalytic activity. In vitro and in vivo characterization confirms that IdeS^M33^ outperforms the wild-type enzyme in the speed and depth of IgG depletion, while maintaining a favorable safety profile. Notably, in a translational application targeting pre-existing immunity, a single dose of IdeS^M33^ effectively neutralized anti-AAV antibodies and restored efficient AAV-mediated hepatic transduction. This study, from rational design and preclinical validation to proof-of-concept application, illustrates the promise of this engineered enzyme in addressing humoral immune barriers.

Our structural analysis, combined with bioinformatic modeling, provides plausible mechanistic insights into the functional enhancement observed for IdeS^M33^. The crystal structure of IdeS^M33^ (residues 47–339) adopts a canonical papain-like cysteine protease fold that is highly conserved with wild-type IdeS.^5^ Structural superposition confirms that the global scaffold remains unperturbed, indicating that the functional gains arise not from large-scale refolding but from local, targeted modifications at the substrate-binding interface.

The most striking local change occurs at position 167. The K167R substitution induces a pronounced side-chain repositioning (∼9.7 Å shift relative to K167 in wild type IdeS), accompanied by a secondary shift of the neighboring Y163 (∼6.9 Å). This reconfiguration reshapes the side-chain microenvironment at a critical region of the binding cleft.^5,20,21^ In the context of the IdeS–IgG Fc complex, R167 is positioned to interact with D265 in the BC loop of the IgG Fc Cγ2 domain.^5^ Given that the BC loop forms part of an extensive recognition interface spanning the hinge region (C229–S239) and the FG loop,^5,20,21^ this reoriented arginine likely enhances electrostatic complementarity and hydrogen-bonding potential, thereby stabilizing the enzyme–substrate complex.

In contrast, the D226E mutation results in a more modest local displacement (∼2.3 Å). While this substitution is conservative in terms of charge, the longer glutamate side chain may improve packing interactions with hinge-region residue A231. Together, these adjustments fine-tune the hydrogen-bonding network across the interface, particularly in regions that are known to stabilize the “open” conformation of the IdeS propeptide-binding loop required for efficient proteolysis.^5,20,21^

Importantly, the structural effects of the M33 mutations are not strictly confined to the immediate substitution sites. Neighboring residues in the Y255–I260 region also exhibit significant shifts (up to 8.6 Å), suggesting that the mutations propagate subtle rearrangements through adjacent loops and surface regions. Such distal but functionally relevant adjustments likely contribute to a more optimal complementarity between IdeS^M33^ and its IgG substrate.

These structural observations align closely with our biophysical data. IdeS^M33^ displays a higher kon and a lower koff, yielding a reduced KD and indicating tighter binding. The enhanced affinity directly translates into superior enzymatic performance: under identical conditions, IdeS^M33^ achieves complete IgG cleavage within 5 minutes—an efficiency not observed with the wild-type enzyme or the individual single mutants.

Collectively, these findings support a model in which discrete but strategically located mutations in IdeS^M33^ stabilize the productive enzyme–substrate complex, lowering the energetic barrier of the rate-limiting step in catalysis. By refining local interfacial geometry and hydrogen-bonding patterns—particularly around R167 and the Y255–I260 region—IdeS^M33^ achieves both tighter binding and faster turnover, offering a structural rationale for its enhanced IgG-cleaving activity.

The enhanced in vitro activity of IdeS^M33^ was robustly translated into superior in vivo efficacy. In rabbits, a species whose IgG is fully cleaved by IdeS and thus represents a highly relevant model,^22^ IdeS^M33^ administration led to a more profound and sustained reduction in serum IgG levels compared to IdeS^WT^ across multiple doses. The lower AUC for serum IgG concentration over time in the IdeS^M33^-treated groups quantitatively confirms its enhanced IgG-depleting capability. The comprehensive pharmacokinetic and pharmacodynamic profiling further defined the therapeutic window within 24 to 48 hours after IdeS administration. While the highest dose (1.8 mg/kg) achieved the greatest drug exposure (AUC) and the most profound IgG nadir, it had the shortest half-life and was associated with the only observed adverse reaction—a significant reduction in red blood cell count. The intermediate dose (0.6 mg/kg) offered the most favorable balance, providing a prolonged half-life and sustained efficacy, whereas the low dose (0.2 mg/kg) remained effective with a modest exposure profile.

The most compelling application demonstrated for IdeS^M33^ is in the field of gene therapy.^23,24^ The presence of nAbs against AAV vectors is a major exclusion criterion for treatment, as it can completely abrogate transduction.^23,24^ Our data show that IdeS^M33^ efficiently degrades both total binding antibodies and neutralizing antibodies against AAV9, reducing nAb titers to negligible levels within 24 hours. This pharmacological intervention had a direct functional consequence: in rabbits with pre-existing anti-AAV9 immunity, a single dose of IdeS^M33^ administered prior to AAV9-eGFP infusion significantly rescued hepatic transduction. The vector genome copy number and the percentage of GFP-positive hepatocytes in the treated group were restored to levels statistically comparable to those in naïve, non-immunized animals. This finding is transformative, as it provides a viable strategy to transiently clear pre-existing humoral immunity, thereby potentially expanding the eligible patient population for AAV-based gene therapies.^25^ The rapid action and specificity of IdeS^M33^ make it an ideal pre-treatment agent, creating a temporary “therapeutic window” for effective vector administration.

Several limitations of this study should be considered. First, the primary in vivo evaluations were conducted in a rabbit model, which, while a relevant species for IgG cleavage by IdeS, may not fully recapitulate the complexity of human adaptive immunity, particularly in terms of antibody repertoires, memory responses, and potential immunogenicity. Second, the study focused primarily on hepatic transduction as a proof of concept; the efficacy of this approach for other target tissues or in the context of different AAV serotypes warrants further investigation.

In conclusion, through structure-guided rational design, we have developed IdeS^M33^, a high-affinity variant of IdeS with enhanced IgG-degrading potency. Its superior efficacy, and favorable safety, validated in a relevant animal model, strongly support its further development. The successful application of IdeS^M33^ in mitigating pre-existing anti-AAV immunity and restoring gene transfer efficiency addresses a critical unmet need, paving the way for its use not only in organ transplantation desensitization and autoimmune disease treatment but also as a key enabling technology for the broader application of in vivo gene therapy. Future work will focus on evaluating its efficacy in primate models and exploring its potential in other clinical scenarios dominated by pathogenic IgG antibodies.

## Materials & Methods

### Homology modeling and structural refinement of the IdeS MG50–Fc complex

Using the crystal structure of the IdeS MG83–Fc complex (PDB ID: 8A47) as a template, the structure of the IdeS MG50–Fc complex was reconstructed via homology modeling using the rosetta.rebuildmodule in PyRosetta (version 4). The modeling procedure employed the ref2015 energy function, incorporating force field optimization and all-atom relaxation. The protocol comprised two steps: first, Cartesian-space FastRelax (repeat number = 5) was performed with Cα positional restraints (weight = 1.0) to eliminate local steric clashes while preserving the template topology; second, 1000 steps of steepest-descent energy minimization were applied for final geometric refinement. The quality of the final model was assessed using ProSA-web, yielding a Z-score below −10, which indicates a structurally reliable model.

### Molecular dynamics simulation

All-atom molecular dynamics (MD) simulations of the IdeS MG50–Fc complex were performed using GROMACS 2026.1 with the Amber19SB force field. The system was solvated in a cubic box (134.6 × 134.6 × 134.6 Å³) containing OPC water molecules and 150 mM NaCl, comprising approximately 77,701 atoms in total. Following energy minimization, the system underwent a 100 ps equilibration phase. Production MD simulations were then conducted for 100 ns in the NPT ensemble at 298.15 K using a velocity-rescale thermostat and a Parrinello-Rahman barostat. Electrostatic interactions were calculated using the Particle Mesh Ewald (PME) method with a 10 Å cutoff, while LINCS constraints were applied to all bonds involving hydrogen atoms, allowing a 2 fs integration timestep.

### Trajectory analysis and identification of key residues

Periodic boundary conditions were corrected using the GROMACS 2026.1 trjconvutility with the-pbc mol-centeroptions. The root-mean-square deviation (RMSD) was calculated for protein backbone atoms, confirming that the simulation reached equilibrium after 20 ns (fluctuation < 0.2 nm). Subsequent analyses were performed on the equilibrated trajectory (20–100 ns) as follows:

(1) Root-mean-square fluctuation (RMSF) of Cα atoms was computed using gmx rmsf;
(2) Hydrogen bonds involving interface residues were analyzed using gmx hbond, applying geometric criteria of a donor–acceptor distance ≤ 0.35 nm and an angle ≤ 30°.

Output metrics included the average number of hydrogen bonds, occupancy (defined as the fraction of simulation time a given hydrogen bond persisted), and the longest continuous absence window. For each residue, we calculated the average number of hydrogen bonds per frame, hydrogen-bond occupancy (the proportion of frames containing at least one hydrogen bond), and the longest continuous window without any hydrogen bond. These metrics were used to assess the stability of the hydrogen-bond network and the potential for transient rupture.

### Expression and purification of IdeS

The codon-optimized gene encoding IdeS was synthesized and cloned into the pGEX-6P-1 vector (GE Healthcare) using BamHI and XhoI restriction sites. Site-directed mutagenesis was performed using a fast mutagenesis kit (Tiangen, Beijing) to generate single and double mutants. All constructs were verified by DNA sequencing. Plasmids were transformed into *Escherichia coli* strain BL21(DE3) cells. Cells were grown in LB medium supplemented with 100 µg/mL ampicillin at 37°C until OD₆₀₀ reached 0.6–0.8, then induced with 0.5 mM isopropyl β-D-1-thiogalactopyranoside at 18°C for 16 h. Cells were harvested, resuspended in PBS, and lysed by high-pressure homogenization. The clarified lysate was loaded onto glutathione Sepharose 4B resin (Cytiva), washed extensively, and the glutathione S-transferase (GST) tag was cleaved on-column using PreScission protease (0.25 mg/mL) at 4°C overnight. The eluted protein was further purified by anion-exchange chromatography on a HiTrap Q column (Cytiva) and size-exclusion chromatography on a Superdex 200 Increase 10/300 GL column (Cytiva) in 10 mM Tris-HCl (pH 8.0) and 100 mM NaCl. Protein purity (>95%) was assessed by SDS-PAGE.

### Crystallization and structure determination

We named the double variant (D226E and K167R) IdeS^M33^. Purified IdeS^M33^ was concentrated to 20 mg/mL in 10 mM Tris-HCl (pH 8.0), 100 mM NaCl. Initial crystallization screening was performed using commercial kits (Hampton Research) via the hanging-drop vapor diffusion method at 18°C. Optimized crystals were grown in 0.1 M Tris-HCl (pH 8.5), 1.0 M sodium citrate. Crystals were cryoprotected in mother liquor supplemented with 10% glycerol and flash-cooled in liquid nitrogen. X-ray diffraction data were collected at the BL01U1 beamline of the Shanghai Synchrotron Radiation Facility (SSRF) on a Pilatus 6M detector at a wavelength of 0.979183 Å. Data were processed and scaled using XDS.^26^

The 2.0 Å crystal structure of IdeS^M33^ (PDB ID 26PT, Extended PDB ID pdb_000026PT) was solved by molecular replacement using Phaser (CCP4 suite) with the IdeS^WT^ structure (PDB 2AU1) as the search model. Automated model building was performed with Phenix AutoBuild Wizard, followed by iterative cycles of manual rebuilding in Coot and refinement with Phenix.refine, including individual B-factor, translation, libration and screw-motion refinement (TLS), and restrained refinement. Water molecules were added during the final stages. Structural validation was performed with MolProbity. Structural analysis and figure generation were done in PyMOL (Schrödinger) and UCSF Chimera. Data processing and refinement statistics are given in Supplementary Table 1.

### Biolayer interferometry

Binding kinetics of IdeS to human IgG were measured by Biolayer interferometry on an Octet Red96 instrument (FortéBio). Biotinylated human IgG (prepared using a biotin quick labeling kit, Frdbbio, ARL0020K) was immobilized onto streptavidin biosensors. Association and dissociation were monitored for 120s and 180s, respectively, with analyte concentrations ranging from 100 to 1600 nM. Data were processed using the Octet Data Analysis Software v9.0 and fitted to a 1:1 binding model to determine the association rate (*k*ₒₙ), dissociation rate (*k*ₒ_ff_), and equilibrium dissociation constant (*K*_D_).

### IgG quantification

Serum IgG concentrations were measured by immunoturbidimetry using a clinical chemistry analyzer. A standard curve was generated using serial dilutions of purified rabbit IgG (0.01–0.64 mg/mL) in PEG-containing buffer. Absorbance at 340 nm was measured after 40 min incubation at 37°C. IgG levels in serum samples were interpolated from the standard curve.

### In vitro IgG cleavage assay

All sera were heat-inactivated at 55°C for 15 min, aliquoted, and stored at –80°C until use. For In vitro serum IgG cleavage assay, 25 μL of serum from human, monkey, canine, and rabbit was treated with 5 μL of IdeS (2 μg/μL) at 37 °C for 1 to 240 min. Aliquots were collected at the indicated time points, denatured in Laemmli buffer, and the cleavage efficiency of the enzyme was analyzed by SDS-PAGE.

### SDS-PAGE and western blot

Samples were resolved on 4–15% gradient polyacrylamide gels under non-reducing conditions and stained with Coomassie Brilliant Blue. To observe the efficiency of IgG cleavage by IdeS, enzymatic cleavage products were separated under reducing SDS-PAGE, transferred to PVDF membranes, and probed with species-specific anti-IgG antibodies: CaptureSelect biotin anti-human IgG-CH1 conjugate (Thermo Fisher, 7103202100) for human and monkey IgG, or anti-rabbit IgG, anti-canine IgG as appropriate. HRP-conjugated secondary antibodies and SuperSignal West Pico Plus substrate were used for chemiluminescent detection (LI-COR Odyssey).

### Animals and ethics

New Zealand White rabbits were procured from Charles River Laboratories (Beijing, China). Animals were housed in a specific pathogen-free facility under controlled conditions (24 ± 2°C, 55% humidity, 12-h light/dark cycle) with free access to food and water. All animal procedures were approved by the Institutional Animal Care and Use Committee of GeneCradle Inc. and conducted in strict compliance with the guideline “Technical specification of ethical review for laboratory animal welfare” (DB11/T 1734-2020).

### Pharmacokinetic and pharmacodynamic analysis

Pharmacokinetic and pharmacodynamic studies were conducted using the rabbit as the animal model. Rabbits were divided into four groups (12 per group) based on dosage: 0 mg/kg (control), 0.2 mg/kg, 0.6 mg/kg, and 1.8 mg/kg. IdeS^M33^ was administered intravenously on Day 0. Changes in blood IgG levels were observed until Day 7. The mean serum IgG level of all rabbits before intravenous administration of IdeS^M33^ is considered as 100%. The IgG level measured at each subsequent time point for each group of rabbits is divided by the mean serum IgG level before IdeS^M33^ injection to obtain the proportion.

To determine pharmacokinetic parameters, rabbit serum samples were mixed with an equal volume of 4 mol/L urea solution and incubated at 25 °C in the dark for 30 min. The mixture was then diluted 20-fold with 1% skim milk solution. The concentration of IdeS^M33^ in rabbit serum was measured using a sandwich enzyme-linked immunosorbent assay (ELISA). Briefly, anti-FabRICATOR Protein G Purified antibody (GENOVIS) was coated onto microplate wells at 200 ng per well and incubated overnight at 2–8 °C. The plate was then blocked with 5% skim milk. IdeS^M33^ treated serum samples and IdeS^M33^ standards (30 ng/ml to 3000 ng/ml) were added and incubated at 37 °C in the dark for 1h. After washing, 100 μL per well of 2 μg/ml His Tag Biotinylated Mouse IgG detection antibody (R&D) was added, followed by incubation at 25 °C in the dark for 1 h. After another wash, 100 μL per well of 1:4000 diluted Streptavidin-HRP secondary antibody (Agilent) was added and incubated at 25 °C in the dark for 1 h. After final washing, 3,3′,5,5′-tetramethylbenzidine substrate was added and incubated at 37 °C in the dark for 10 min. The reaction was stopped with 100 μL per well of stop solution. Absorbance was measured at 450 nm and 630 nm using a microplate reader. IdeS^M33^ concentrations in samples were determined based on a standard curve fitted from the standard absorbance readings.

### AAV9 vector production and purification

Wild-type AAV9 (AAV9^WT^) and a recombinant vector encoding enhanced green fluorescent protein (rAAV9-eGFP) were produced using a baculovirus-based Bac-to-AAV system followed by cesium chloride (CsCl) gradient ultracentrifugation. Briefly, recombinant baculoviruses encoding AAV9 Rep/Cap genes and the transgene expression cassette flanked by inverted terminal repeats were co-transfected into Sf9 cells at a multiplicity of infection of 0.1. Seventy-two hours post-infection, cell lysates were clarified by centrifugation, and vectors were purified using an AAV9-specific affinity column (AVB Sepharose, Cytiva). Bound virions were eluted with 100 mM glycine and 500 mM arginine (pH 2.5), immediately neutralized, and subjected to CsCl density gradient ultracentrifugation (65,000 rpm, 15°C, 20 h) in a 70 Ti rotor (Beckman Coulter) to isolate full capsids. Purified vectors were buffer-exchanged into sterile PBS containing 0.001% Pluronic F-68, filtered through a 0.22-µm membrane, and stored at –80°C.

### Anti-AAV IgG cleavage assay

After mixing empty capsids of AAV serotype9 and aluminum adjuvant at a volume ratio of 1:1, Rabbits were subcutaneously immunized at least at 5 sites on the back at a dose of 1.5 mg per rabbit. A second immunization was performed using the same method and dose two weeks later. Three weeks after the second immunization, IdeS^M33^ (0.2 mg/kg) or solvent was administered.

To quantify the cleavage capacity of IdeS^M33^ against AAV binding IgG, the capsid protein antigen of AAV serotype9 was coated onto microplate wells at 100 ng/well and incubated at 2-8°C overnight. Wells were then blocked with 5% skim milk. Serially diluted test serum samples (starting dilution 1:100, followed by 2-fold serial dilutions) and negative controls were added and incubated at 37°C for 1 hour. After washing, horseradish peroxidase (HRP)-conjugated Goat Anti-Rabbit IgG secondary antibody (abbkine) was added at 50 µL/well and incubated at 37°C protected from light for 30 min to 1 h. Following another wash, 3,3′,5,5′-tetramethylbenzidine was added for color development at room temperature protected from light for 5-15 minutes. The reaction was then stopped with stop solution. Absorbance (OD) was measured at 450 nm (primary wavelength) and 630 nm (reference wavelength) using a microplate reader. The antibody titer was defined as the maximum dilution factor at which the OD value reached 2.1 times that of the negative control.

### AAV hepatic transduction with pre-existing anti-AAV antibodies treated with IdeS^M33^

To investigate whether IdeS^M33^ treatment could enhance the efficiency of subsequent AAV9 re-administration following initial immunization, rabbits were treated with or without 0.2 mg/kg IdeS^SM33^ for one day, two months after the initial immunization. Subsequently, all rabbits received an intravenous injection of rAAV9-eGFP (2 × 10¹² vg/animal) via the tail vein. To determine whether IdeS^M33^ could effectively mitigate the impact of AAV9 neutralizing antibodies on AAV9 transduction efficiency, GFP content in liver tissue was quantified by quantitative PCR and liver histopathology.

### Statistical Analysis

Data are presented as mean ± standard deviation. The area under the curve (AUC) of antibody concentration over time was used to determine if there were significant differences in the cleavage efficiency of IdeS^WT^ versus the IdeS^M33^ mutant. To evaluate the time-dependent cleavage effect of IdeS on IgG, a one-way ANOVA was performed to assess the statistical significance of IgG reduction across time points. To evaluate the safety of IdeS^M33^, a two-way ANOVA was used to analyze the effects of dose (0.0, 0.2, 0.6, and 1.8 mg/kg) and time, as well as their interaction, on the safety indicators. To determine whether IdeS^M33^ can improve AAV transduction efficiency in the presence of pre-existing immunity, a t-test was used to compare AAV-mediated gene transduction and expression efficiency between (1) rabbits with pre-existing immunity that either received or did not receive IdeS^M33^ treatment, and (2) rabbits with pre-existing immunity treated with IdeS^M33^ versus rabbits without pre-existing immunity. All statistical analyses were performed using GraphPad Prism (version 10). A two-tailed P value of < 0.05 was considered statistically significant.

## Data and code availability

All data reported in this manuscript will be shared with the lead contact upon request.

## Supporting information

supplemental files in one PDF file

## Acknowledgments

This work was partially supported by the Beijing E-Town Cooperation and Development Foundation (grant number YJXJ-JZ-2021-0014-08).

## Author contributions

X.D., C.C., W.M. and X.W. conceived and supervised this study. W.M. designed the experiments, which were performed by K.Z., Z.W., Z.R., Co.C., Y.X., D.H., Z.Y., H.N., J.Q., P.G., W.Y., Y.D., X.L., and Z.D. The paper was written by Y.W., W.M., and C.C.

## Supplemental information (1)

