## supplemental files in one PDF file for "An engineered IdeS variant with enhanced activity and performance for IgG degradation"

### Supplementary materials

**Supplementary Figure 1. LigPlot analysis of the hydrogen-bond interaction network at the IdeS MG50–IgG-Fc interface.** IdeS MG50 (chain C) interacts with IgG-Fc (chains A and B) through two distinct hydrogen-bond networks. Residues involved in the B–C interface are E132, K167, R185, T189, and R204 (A); those participating in the A–C interface are E66, D67, K84, A94, D226, K228, V258, R259, A310, and G319 (B).

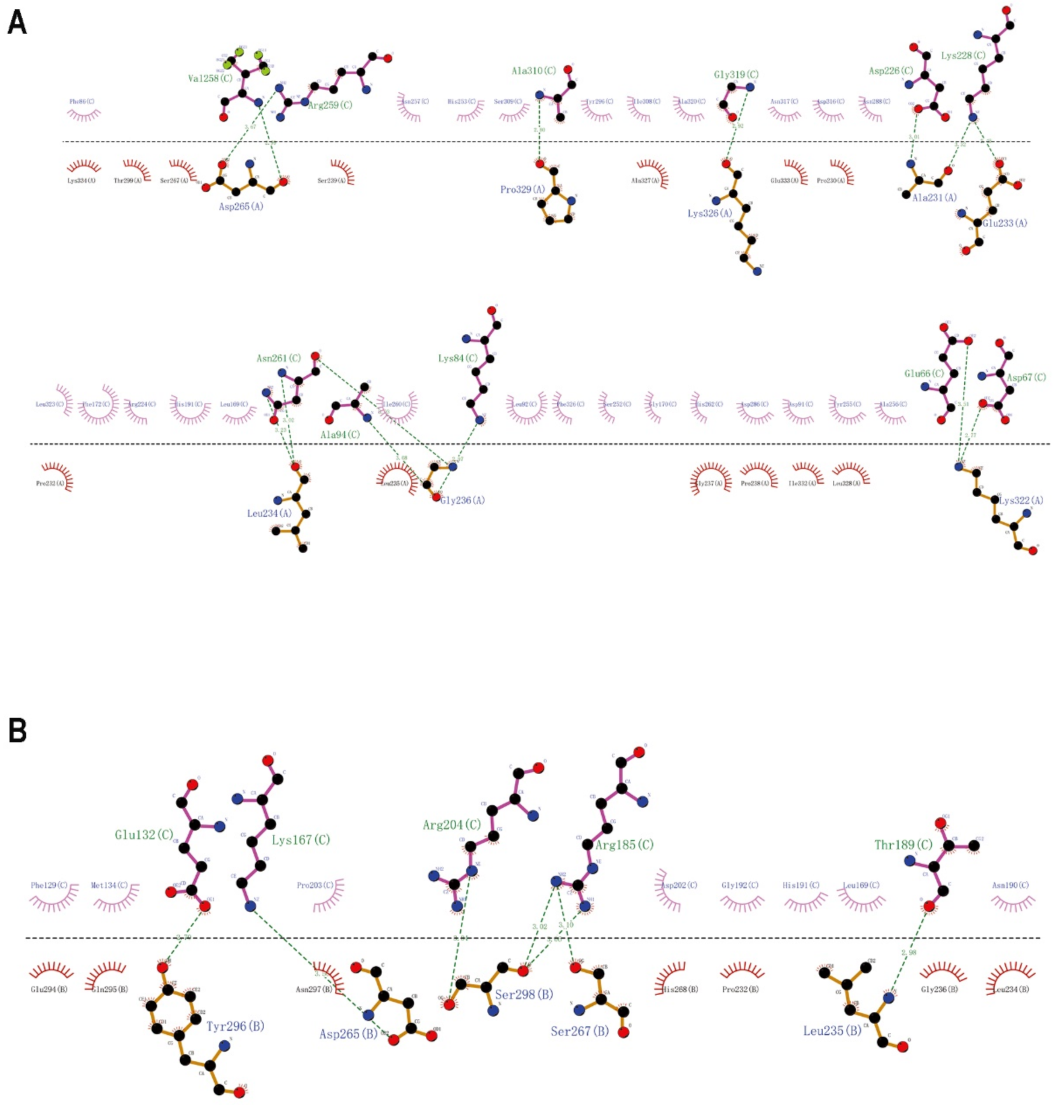

**Supplementary Figure 2. Hydrogen-bond dynamics at the B–C and A–C interfaces of the IdeS MG50–IgG-Fc complex.** Analysis of hydrogen-bond formation and stability between IdeS MG50 (chain C) and IgG-Fc (chains A and B), highlighting dynamic behavior at the B–C and A–C interfaces.

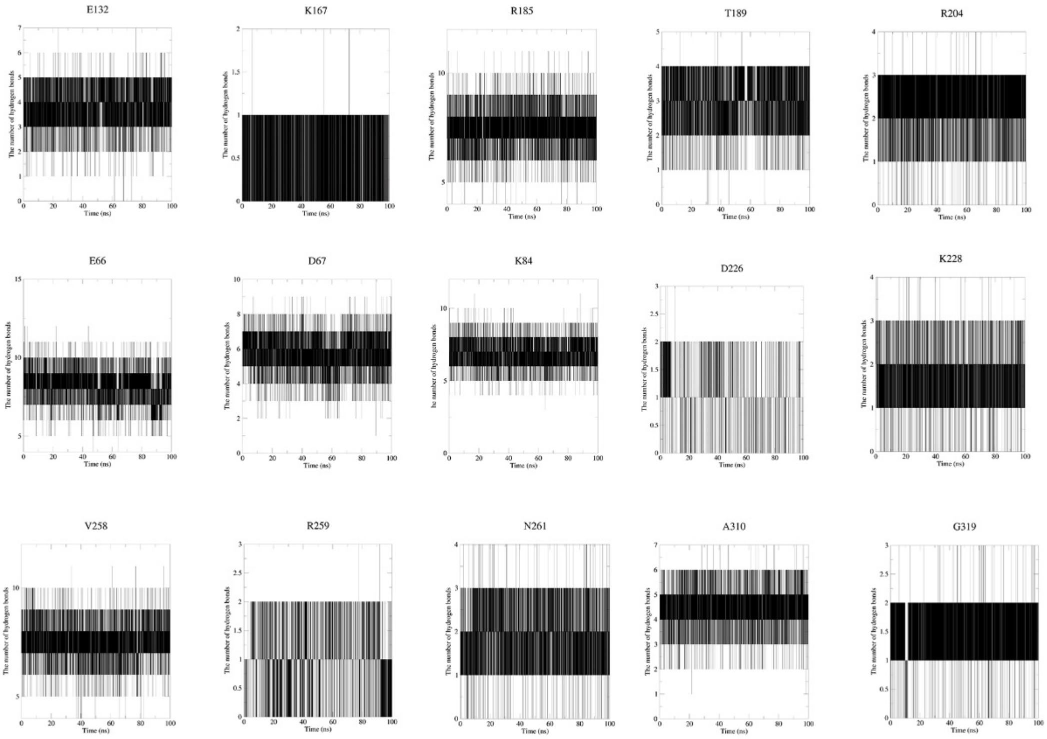


| Residues | Average H-bonds | H-bond occupancy, % | Longest H-bond absence window, ps |
|---|---|---|---|
| E132 | 3.86 | 99.95 | 10 |
| K167 | 0.71 | 70.65 | 90 |
| R185 | 7.50 | 100 | 0 |
| T189 | 3.02 | 99.94 | 10 |
| R204 | 2.37 | 99.25 | 10 |
| E66 | 8.15 | 100 | 0 |
| D67 | 5.70 | 100 | 0 |
| K84 | 6.84 | 100 | 0 |
| D226 | 1.01 | 96.38 | 30 |
| K228 | 1.77 | 97.6 | 20 |
| V258 | 7.53 | 100 | 0 |
| R259 | 0.96 | 85.21 | 260 |
| N261 | 1.91 | 98.85 | 20 |
| A310 | 4.52 | 100 | 0 |
| G319 | 1.51 | 98.64 | 20 |

**Supplementary Figure 3. Safety assessment of IdeSM33 in rabbits.** To evaluate the safety of IdeSM33, rabbits were administered intravenous injections of IdeSM33 at doses of 0.0, 0.2, 0.6, and 1.8 mg/kg. Body temperature was recorded for 8 days (A), and body weight was monitored for 7 weeks (B: male rabbits; C: female rabbits). Complete blood counts were examined weekly up to the third week (D: male rabbits; E: female rabbits). Blood biochemistry was analyzed before injection and on day 17 post-injection (F: male rabbits; G: female rabbits). The P-values shown in the figure represent the interaction between the time factor and the dose group in a two-way ANOVA. WBC, white blood cell; LYM, lymphocyte; MONO, monocyte; NEUT, neutrophil; PLT, platelet; RBC, red blood cell; ALT, alanine aminotransferase; AST, aspartate aminotransferase; CK, creatine kinase; CRE, creatinine.

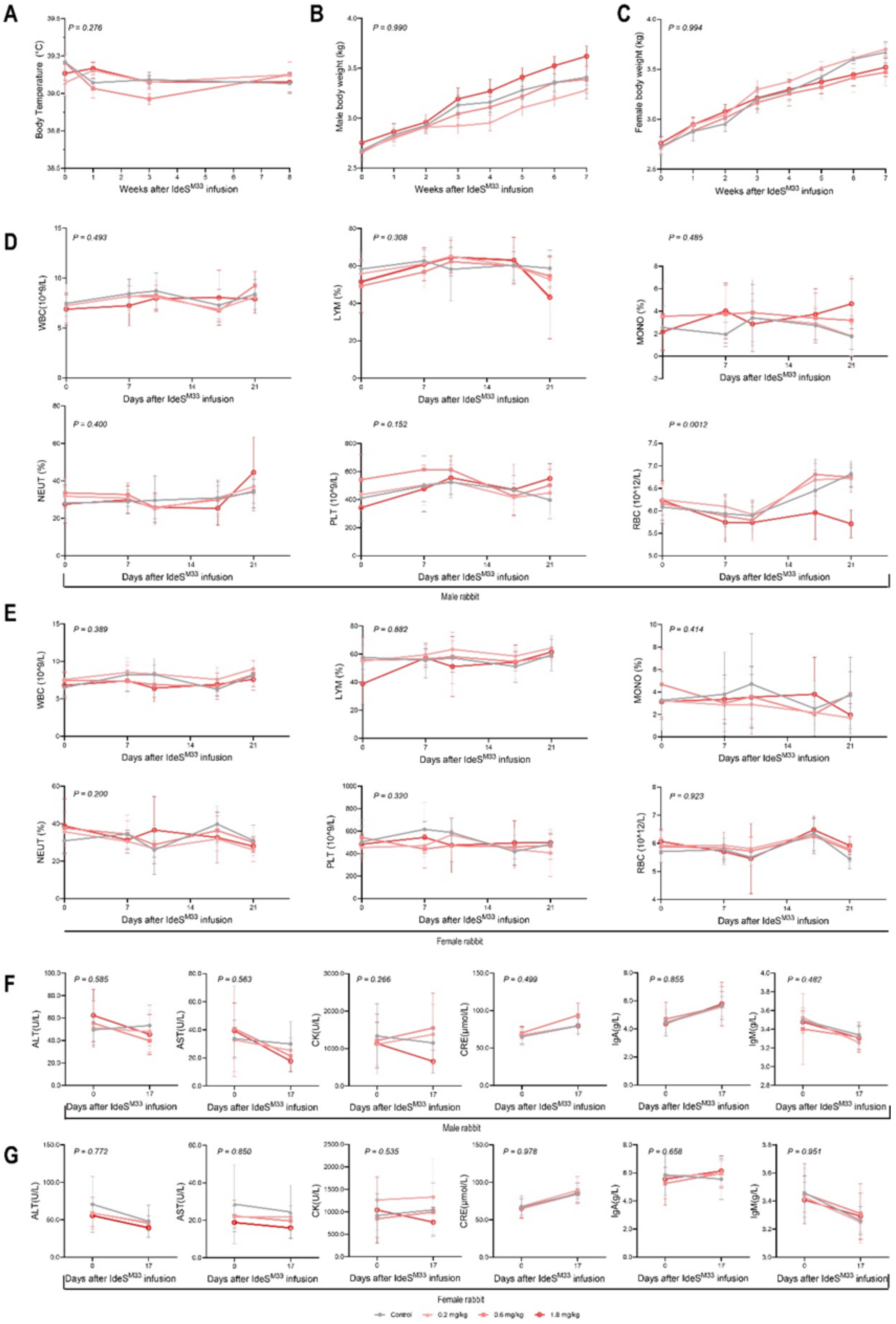
A
P = 0.276
Body Temperature (°C)
Weeks after IdeS$^{M33}$ infusion
B
P = 0.990
Male body weight (kg)
Weeks after IdeS$^{M33}$ infusion
C
P = 0.994
Female body weight (kg)
Weeks after IdeS$^{M33}$ infusion
D
P = 0.493
WBC(10^9/L)
P = 0.308
LYM (%)
P = 0.485
MONO (%)
P = 0.400
NEUT (%)
P = 0.152
PLT (10^9/L)
P = 0.0012
RBC (10^12/L)
Days after IdeS$^{M33}$ infusion
Male rabbit
E
P = 0.389
WBC (10^9/L)
P = 0.882
LYM (%)
P = 0.414
MONO (%)
P = 0.200
NEUT (%)
P = 0.320
PLT (10^9/L)
P = 0.923
RBC (10^12/L)
Days after IdeS$^{M33}$ infusion
Female rabbit
F
P = 0.585
ALT(U/L)
P = 0.563
AST(U/L)
P = 0.266
CK(U/L)
P = 0.499
CRE(μmol/L)
P = 0.855
IgA(g/L)
P = 0.482
IgM(g/L)
Days after IdeS$^{M33}$ infusion
Male rabbit
G
P = 0.772
ALT(U/L)
P = 0.850
AST(U/L)
P = 0.535
CK(U/L)
P = 0.978
CRE(μmol/L)
P = 0.658
IgA(g/L)
P = 0.951
IgM(g/L)
Days after IdeS$^{M33}$ infusion
Female rabbit
Control
0.2 mg/kg
0.6 mg/kg
1.8 mg/kg

Supplementary Table 1. Data collection and refinement statistics.

| | IdeS$^{m33}$ |
|---|---|
| **Data Collection** | |
| X-ray Source | SSRF BL02U1 |
| Wavelength (Å) | 0.979183 |
| Space group | $P2_12_12$ |
| Unit cell parameters (Å; °) | a=64.3, b=87.7, c=56.4; α=β=γ=90.0 |
| Resolution range (Å) | 50.0-1.91 (1.96-1.91)* |
| Unique reflections | 25,168 (1,586) |
| Completeness (%) | 99.0 (88.0) |
| Redundancy | 11.6 (6.8) |
| I/σ(I) | 27.4 (4.1) |
| $R_{merge}$ (%) | 5.6 (63.5) |
| $R_{pim}$ (%) | 1.7 (25.1) |
| $R_{meas}$ (%) | 5.9 (68.7) |
| $CC_{1/2}$ | 0.999 (0.803) |
| **Refinement** | |
| Resolution range (Å) | 30.49-1.91 (1.978-1.91) |
| Reflections used in refinement | 25,064 (2,197) |
| Reflections used for R-free | 1,175 (111) |
| $R_{work}$ (%) | 18.7 (28.7) |
| $R_{free}$ (%) | 21.10 (33.8) |
| Number of non-hydrogen atoms | 2,474 |
| Protein | 2,333 |
| Solvent | 141 |
| Average B-factors | 45.0 |
| Protein | 45.0 |
| Solvent | 44.8 |
| r.m.s. deviations | |
| Bond lengths (Å) | 0.006 |
| Bond angles (°) | 0.85 |
| Ramachandran | |
| Favored (%) | 97.2 |
| Allowed (%) | 2.8 |
| Outliers (%) | 0.0 |
